# Decomposing a San Francisco Estuary microbiome using long read metagenomics reveals species and species- and strain-level dominance from picoeukaryotes to viruses

**DOI:** 10.1101/2023.06.30.547288

**Authors:** Lauren M. Lui, Torben N. Nielsen

## Abstract

Although long read sequencing has enabled obtaining high-quality and complete prokaryotic genomes from metagenomes, many challenges still remain to completely decompose a metagenome into its constituent genomes. These challenges include obtaining enough biomass, high-molecular weight DNA extraction, determining the appropriate depth of sequencing, and bioinformatics challenges to separate closely related genomes. This study focuses on decomposing an estuarine water metagenome from USGS Station 36 in the South San Francisco Bay into its constituent genomes and counting the number of organisms present. To achieve this, we developed a new bead-based DNA extraction method, a novel bin refinement method, and sequenced the sample with 150 Gbases of nanopore sequencing. With our results, we were able to estimate that there are ∼500 bacteria and archaeal species in our sample, obtain 68 high-quality bins (>90% complete, <5% contamination, ≤5 contigs, no contigs shorter than 100 Kbases, and all ribosomal and necessary tRNA genes). Since we pre-filtered the sample at 11μm and then collected directly on to a 0.1μm filter, we also obtained many contigs of picoeukaryotes, environmental DNA of larger eukaryotes such as mammals, complete mitochondrial and chloroplast genomes, and detected ∼40,000 viral populations. This deep analysis of the taxonomy of the sample down to the strain and individual contig level allowed us to find that among picoeukaryotes, prokaryotes, and viruses there are likely only a few strains that comprise most of the species abundances. These results also indicate that to truly decompose a metagenome into its constituent genomes, we likely need 1Tbase of sequencing.

If you are reading this preprint, know that this is the paper we wanted to write, but it will likely be shortened for submission to a journal.

## Introduction

The effect of climate change on microorganisms and their potential to positively or negatively affect climate change is of growing importance (Cavicchioli et al. 2019; Brennan and Logares 2023). Ocean and estuarine microbiomes are of major interest because of their contributions to greenhouse gas production and consumption, as well as their fundamental roles in global element cycling (Brennan and Logares 2023; Cavicchioli et al. 2019). For example, even though the ratio of the total biomass of marine phytoplankton to that of terrestrial flora is approximately 1:100, marine microbial communities contribute approximately half of net primary production (*e*.*g*., photosynthesis) of the Earth (Behrenfeld 2014; Field et al. 1998). Despite the importance of modeling microbial communities to predict the effects of climate change, a fundamental unit for helping us study environmental microbiomes still lies tantalizingly out of reach: complete genomes from unculturable microbes.

Metagenomics has provided an easy, cost-effective way to study the diversity, community structure, functional potential, and behavior of environmental microbial communities, especially since they require a microscope to be seen (unless there is a phytoplankton bloom and then they can be seen from space (Behrenfeld 2014)) and many species cannot be cultured in the laboratory. Likely most modern microbiologists that use sequencing in their studies know the genomes and phylogenetic placement of their subjects of study better than the morphology. However, despite this focus on genomes, obtaining complete bacterial and archaeal genomes from metagenomes is still a difficult task (Lui et al. 2021; Chen et al. 2020), especially if there is high strain diversity. This is evidenced by the low number of genomes in databases in comparison to estimated microbial diversity (tens of thousands of genomes compared to millions of species) (Albertsen 2023) and that the standard for a “medium quality” metagenome assembled genome (MAG) is 50% completeness and 10% contamination (Albertsen 2023; Bowers et al. 2017). Incomplete genomes can interfere with understanding a species’ full functional capabilities (Eisenhofer et al. 2023), have contaminating genes from other genomes (Albertsen 2023), and make accurate estimates of species abundance in samples difficult. Beyond prokaryotic genomes, viral and eukaryotic metagenomics are also challenging but important for understanding the dynamics of the entire marine microbial community (Gregory et al. 2019; Patin and Goodwin 2022; Sunagawa et al. 2020).

In the last 5 years, long read sequencing (as developed by Oxford Nanopore Technologies and Pacific Biosciences) has facilitated the assembly of complete or nearly complete genomes from metagenomes from a variety of environments (Cuscó et al. 2021; Singleton et al. 2021; Overholt et al. 2020; Moss et al. 2020; Maghini et al. 2021; Arumugam et al. 2021; Stewart et al. 2019; Kim et al. 2022; Patin and Goodwin 2022), as well as plasmids and viruses (Beaulaurier et al. 2018, 2020; Dai et al. 2022; Martin et al. 2021), although the data comes with its own technical and bioinformatics challenges. While it is known that high-quality, high-molecular-weight (HMW) DNA is needed for the best results from long read sequencing (Maghini et al 2021), many groups have been using standard DNA extraction methods as input to their long read library preparation and this has often resulted in suboptimal read length and quality. For environmental microbiomes, relic DNA is also an issue because not only does it represent dead organisms, the DNA fragments tend to be short (<500 bp) (Lennon et al. 2018). Although longer reads can help overcome repeats that fragment assemblies (Cahill et al. 2010; Lui et al. 2021), binning algorithms are generally not designed to handle the longer contigs and may fail to bin properly in part due to their reliance on accurate coverage measures (He et al. 2015; Jégousse et al. 2021). Effectively using long read sequencing to obtain complete genomes from metagenomes still needs to overcome the challenges of obtaining enough biomass, high-quality high-molecular weight DNA extraction, and bioinformatics challenges.

In this study, we sought to establish methods to obtain complete or high-quality microbial genomes to study and model the microbiome of the San Francisco Estuary (SFE), the largest estuary on the west coast of the United States. The SFE has high nutrient loadings, especially nitrogen and phosphorus, that are higher than other estuaries already impaired by eutrophication syndrome (Cloern et al. 2020). This along with increased algal toxins and primary production in recent years supports the hypothesis that the SFE is at a critical tipping point and better models are urgently needed to investigate the health of the ecosystem. We sought to address the following methodological questions:

- Can we optimize DNA extraction from filters for long read sequencing of marine samples?
- What depth of nanopore sequencing is necessary to obtain complete genomes of all prokaryotic species in a sample? Related, is it necessary to separate size fractions of eukaryotes, prokaryotes, and viruses to enrich for sequencing?
- Can we improve bioinformatics methods to assist with obtaining complete genomes from nanopore sequencing and can we separate strains?

Here we present a first look at deep Nanopore sequencing of a sample from the South San Francisco Bay (Figure 1A). We have sequenced this estuarine microbiome sample with 150 GBases of nanopore sequencing, which is deeper long read sequencing than any other marine or estuarine study that we know of and is deeper than most other long read microbiome studies. We developed our own filtering and HMW DNA extraction protocols to improve the quality and amount of DNA available for sequencing (Figure 1B), especially to have enough for size selection. Using a novel binning method (Figure 1C), we were able to complete or nearly complete 68 microbial genomes, as well as identify ∼1900 plasmids and ∼40,000 viral populations. We define near complete microbial genomes as greater than 90% complete, less than 5% contamination, no more than 5 contigs and no contigs shorter than 100 Kbases as well as having full ribosomal complements and all necessary tRNAs. This definition extends the high-quality draft definition previously published (Bowers et al. 2017). In addition to the bacterial and archaeal members, we also generated partial genomes for several eukaryotes and we obtained multiple full length mitochondrial and chloroplast genomes; these will be the focus of later studies.

**Figure 1:**
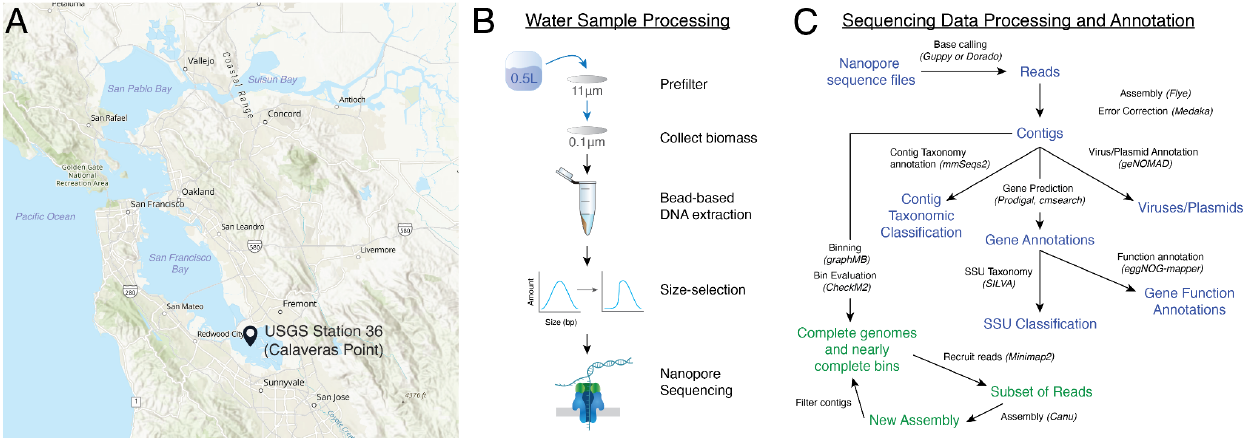
Overview of Sample and Data Processing. (A) Map of the San Francisco Estuary and the location of USGS Station 36. Background map created using ArcGIS® software by Esri with the baseline map “World Street Map (with Relief) - no labels” (ID:5514df296eb348aeac7bf0006663cf48). (B) Overview of water sample processing. (C) Overview of data processing of nanopore sequencing data (blue) and iterative binning method (green).

Although we were not able to complete all of the genomes in our sample, we gained valuable information and determined that we have likely obtained most of the diversity in the sample and genome bins for most species present, approximately 40% of our sample consists of viruses, and we can estimate the depth of sequencing needed to separate strains and complete all of the genomes in our samples. These results represent advances in using long reads to obtain microbial genomes from metagenomes, and an initial look into our project to decompose the entire SFE microbiome.

## Materials and Methods

### Water Sampling and Processing

Ten liters of surface water was collected by Erica Nejad and the crew of the USGS R/V David H. Peterson on July 20, 2022 from USGS Station 36 (Figure 1A). We received the water the same day and immediately transported it to a cold room kept at 4°C. Prefiltering through an 11 μm cellulose filter (Zenpore qualitative grade 1, ST001-125) to remove particulate matter and eukaryotes was done within the same day. Final filtering through 0.1 μm PES filters (Pall, 60311) was carried out within 24 hours. We passed 0.5 L of water through each of twenty 0.1 μm filters and immediately transferred them to a -20°C freezer.

### DNA Extraction and Size Selection

We extracted 13.7 μg of high molecular weight (HMW) DNA from four filters using a gentle enzymatic lysis followed by a bead-based cleanup. The extracted DNA was further size-selected using a Circulomics Short Read Eliminator XS kit (deplete DNA fragments <10kb) yielding a total of 4.1 μg of size-selected DNA. The quality of the DNA was checked on a Nanodrop to ensure that the OD_600_ 260/230 and 260/280 ratios were >1.8 and the DNA length distribution was checked using a Femto Pulse (Agilent Technologies; Supplementary Figure 1).

### Nanopore Library Preparation and Sequencing

Approximately 1μg of size-selected DNA was used as input to each of four library preps. We generally followed published Oxford Nanopore Technology protocols but extended incubation times. First, we did the DNA repair and end prep using New England Biolabs NEBNext FFPE DNA Repair Mix and NEBNext Ultra II End repair / dA-tailing Module. The reactions were 48μL of size-selected DNA (DNA CS was omitted), 3.5μL of NEBNext FFPE DNA Repair Buffer, 2μL of NEBNext FFPE DNA Repair Mix, 3.5μL of Ultra II End-prep reaction buffer, and 3 μl Ultra II End-prep enzyme mix. The reaction was carried out in a thermocycler for 25 minutes at 20°C followed by 5 minutes at 65°C. The reaction was cleaned up by using Ampure XP beads at a 1:1 ratio (60 μL) and with two 70% ethanol washes. To elute the DNA, the beads were incubated in water for 10 minutes at room temperature.

Next, the library was prepared using the SQK-LSK110 kit for R9.4 flow cells or SQK-LSK112 kit for R10.4 flow cells. The adapter ligation reaction was extended to 30 minutes. The reaction was cleaned up by using Ampure XP beads at a 2:3 ratio (40 μL) and washed twice with the supplied Long Fragment Buffer to enrich for DNA fragments longer than 3kb. DNA was eluted from beads with 16uL of Elution Buffer and incubated at 37C for 10 minutes. The library was run on a Femto Pulse to confirm library quality (Supplementary Figure 2). Oxford Nanopore Technologies (ONT) long read sequencing was done using two R9.4.3 MinION flow cells and two R10.4 PromethION flow cells. All runs were set to 72 hours and allowed to go to completion.

### Basecalling, Polishing, and Assembly of Nanopore Data

Basecalling was performed using the Guppy version 6.2.1+6588110a6. The MinION runs used the r9.4.1_450bps_sup model and the PromethION ones used the r10.4_450bps_hac model. We obtained a total of 149,898,872,235 bases (150 Gbases) that passed default Guppy quality control. Read N50/N90 was 11,935 / 3,570 bases.

The long read assembly was done using Flye version 2.9.1-b1780. Other than specifying metagenomic mode, all parameters were left at their default settings. Flye selected a minimum overlap of 4,000 bases. The total length of the assembly is 2,948,820,500 bases (2.95 Gbases), there were 107,977 contigs, the N50 was 62,203 bases, the longest contig was 3,483,052 bases (3.5 Mbases) and the average coverage was 36X.

The long read assembly was error corrected using Medaka version 1.7.1 with model r104_e81_hac_g5015. We selected the model per ONT recommendations based on the available models.

### Illumina Metagenomics Sequencing and Assembly

In addition to the long read sequencing, we also generated 21 Gbases of 2 × 150 bp reads using Illumina sequencing using DNA from the same extraction. DNA was slightly sheared by pipetting up and down 10 times and was sent to Novogene Corporation Inc. (California, USA) for library prep and 2×150bp sequencing on a NovaSeq6000 (Illumina, USA). The run resulted in 21Gb of data with 140,278,770 raw reads, 92.2% reads with quality scores >Q30. The Illumina reads were trimmed and filtered using BBtools version 38.96 as described in (Lui et al. 2021).

The Illumina reads were assembled using SPAdes version 3.15.5 (Prjibelski et al. 2020). The total length of the assembly is 948,090,524 bases (0.95 Gbases), there were 813,394 contigs, the N50 was 1,270 bases, the longest contig was 194,182 bases and the average coverage was 22.15X. These numbers were calculated using Quast (Mikheenko et al. 2016) with default parameters.

### Protein and RNA Gene Annotation

Gene calling was done using Prodigal version 2.6.3 (Hyatt et al. 2010) for both Nanopore and Illumina assemblies. For both datasets, Prodigal was set to not call genes across edges of contigs, to force a full motif scan and to use metagenomic mode. For the Nanopore assembly, Prodigal found 4,153,507 genes. For the Illumina assembly, Prodigal found 5,492,197 genes. Both Nanopore and Illumina assemblies were annotated against the full Rfam database using the Infernal package (Nawrocki and Eddy 2013) with default parameters. Taxonomy of SSUs was done using the SILVA website and manual comparison with BLAST searches.

### Taxonomy Assignment of Contigs

We used MMSeqs2 (Mirdita et al. 2021) with default parameters to assign taxonomy to the Nanopore contigs. Out of the total 107,977 contigs, 95,282 were assigned a taxonomy and 22,761 were assigned down to the species level. We also used Kaiju (Menzel et al. 2016) with default parameters and the db_nr_euk database.

### Taxonomy Assignment and Completeness Check for Bins

We used GTDB-Tk v2 (Chaumeil et al. 2022) for assigning taxonomy to both complete genomes and bins. For completeness determination, we used CheckM2 (Chklovski et al. 2022). In both cases, we ran with default parameters.

### Plasmid and Virus Classification and Analysis

We used geNomad (Camargo et al. 2023) with default parameters to classify contigs as plasmids and viruses. We determined viral populations as sequences that had ≥95% average nucleotide identity (ANI) as defined by (Gregory et al. 2019). We used *dedupe*.*sh* from BBtools (BBMap – Bushnell B. – sourceforge.net/projects/bbmap/) using the parameter minidentity=95. Sequences that were reported as duplicates and contained sequences were removed from the total number of sequences to get the number of viral populations.

### Tree Building and Visualization

All trees were built using IQTree (Nguyen et al. 2015) with the model set to GTR+I+G and the number of fast bootstraps to 3,000. The trees we are dealing with are relatively small and we did not see a need for model finding. iTOL (Letunic and Bork 2021) was used for tree visualization.

### Data Deposition

We will deposit all refined bins that meet the requirements of our near complete genome definition. We have experienced too many issues with poorer quality bins in various repositories and we do not wish to contribute to the problem. We are currently working on getting NCBI Accession numbers and these will be included in the final manuscript.

## Results and Discussion

### Processing and Sequencing of Water Samples from USGS Station 36

We received water samples from USGS from Station 36 in the South San Francisco Bay (Figure 1A). Station 36 is one of multiple USGS monitoring stations in San Francisco Bay and the sampling methods are described in (Schraga and Cloern 2017). Salinity at the station was 32.70 Practical Salinity Units (PSUs) and the temperature was 21.47 °C.

To improve collection of the entire microbial population (<11μm), we chose to collect biomass directly onto a 0.1μm filter after prefiltering (Figure 1B), rather than sequential filtering that is done in many other water metagenomics studies (Sunagawa et al. 2020). This reduces possible loss from sequential filtering and allows us to capture ultra-small bacteria and viruses (Luef et al. 2015). To extract high-quality HMW DNA for long read sequencing, we used an extraction method we developed. It uses gentle enzymatic lysis and carboxyl beads to reduce DNA shearing and maximize DNA quality (OD_600_ 260/230 and 260/280 > 1.8).

We initially sequenced nanopore libraries on two R9.4.1 flow cells, but found that we wanted deeper sequencing to improve coverage of some low abundance species and thus improve their assemblies. We obtained additional sequencing with two R10.4 PromethION flow cells, for a total of ∼150Gbases of data. We obtained a read N50 of 11,935bp. That is about twice that of other long read metagenomes of environmental water samples, which is typically 1000-6000 bp (Overholt et al. 2020; Haro-Moreno et al. 2021; Patin and Goodwin 2022). We also obtained 20 Gbases of Illumina sequencing from the same DNA used for the nanopore libraries.

### Taxonomic Composition of the Sample at the Superkingdom Level

Although the main goal of this study was to assemble high-quality bacterial and archaeal genomes, we also attempted to classify all 107,977 contigs in the sample at least to the superkingdom (domain) level. This effort also provided an overview of the organisms that we captured on our filters (0.1-11μm), and revealed the amount of environmental DNA we could detect from larger organisms, from microplankton to sea mammals. Theoretically, the longer contigs afforded by the nanopore sequencing provide more sequence and gene resolution on which to base classification.

We used three classification methods in conjunction with manual curation to analyze the contigs in our samples. First, we used geNomad, a deep neural network method, to classify contigs as mobile genetic elements (*e*.*g*., plasmids and viruses) and to assign viral taxonomy. The authors of geNomad did extensive curation of their training data from genomes, plasmids, and viruses. They also added in viral data from specific papers to augment the training data for viral sequences. Second, we used MMseqs2, which assigns taxonomy based on protein genes, to classify contigs. GTDB was used as the reference database, so this classification was biased towards bacteria and archaea. Third, we used Kaiju to classify contigs. We used the Kaiju database that contains a subset of the NCBI BLAST nr database containing all proteins from archaea, bacteria, viruses, fungal, and microeukaryotic reference genomes (db_nr_euk). Theoretically this method should be the best at assigning eukaryotic classifications to the contigs. Finally, we manually curated contigs as either mitochondria or chloroplasts by using NCBI BLAST (Camacho et al. 2009) on the SSUs.

Not unexpectedly, using multiple classification methods leads to discrepancies, but combining all of the results led to the conclusion that of the contigs we assembled, 4% are eukaryotic, 0.3-0.7% are archaeal, 38% are bacterial, 40% are viral, and 14% are unclassified/ambiguous (Figure 2A). If we were to more conservatively assign contigs as viral, the percentage of viruses would go down to 25% and the number of unclassified contigs would increase to 29%. We encourage the reader to see the Supplementary Methods for full results from each of the classification methods, assumptions, and reasoning for reaching our final classification of the contigs. The unclassified/ambiguous contigs had a median length of ∼5000 bp; possibly many of these contigs could not be classified because they were too short, rather than the databases not having representative sequences. Sorting through the different classification methods also points out the hazards of relying on one method for classification, even if the focus of a study is purely prokaryotic, viral, or eukaryotic.

**Figure 2:**
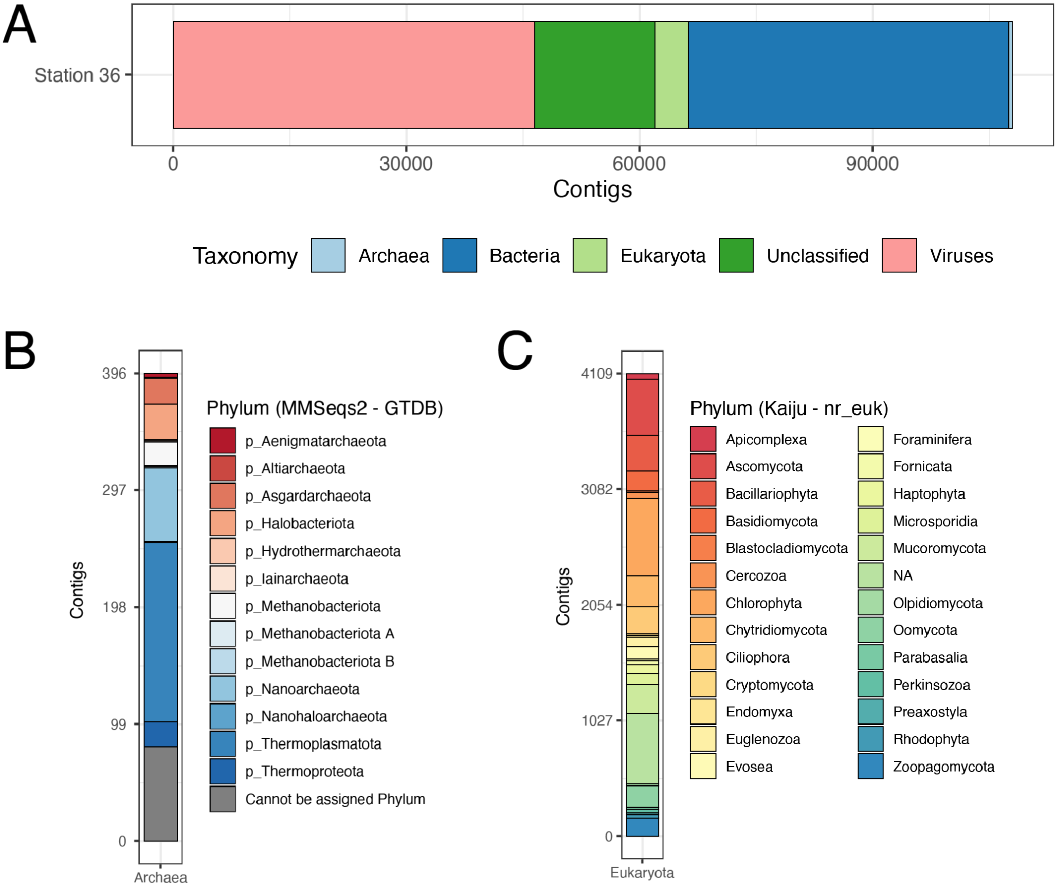
Contig Classification. (A) Curated taxonomy at the superkingdom level based on geNomad, Kaiju, and MMseqs2 (Supplementary Methods). (B) Archaeal contig classification at the phylum level based on MMseqs2. (C) Eukaryotic contig classification at the phylum level based on Kaiju using the nr_euk database. Note that these results only include microeukaryotes and fungi (limitation due to Kaiju database). The eukaryotic contigs classified based on SSU analysis are not shown here because a different classification database was used (SILVA or manual curation based on NCBI nr database search).

### Bacterial taxa comprise nearly all prokaryotic contigs, while archaeal taxa are rare

Unsurprisingly, given that we prefiltered the water at 11μm, archaea and bacteria composed a large proportion of the contigs. There were 40,986 bacterial contigs (38%) and we discuss the bacterial populations in further detail later in the manuscript. We only obtained one contig with a complete archaeal SSU, which was classified as in the *Nitrosopumilus* genus by both MMSeqs2 and SILVA (Supplementary Tables 1 and 2). This is consistent with a recent 16S study where a *Nitrosopumilus*-like operational taxonomic unit (OTU) was the only dominant archaeal OTU in the South Bay of the SFE (Rasmussen et al. 2021). In a subsequent study, the authors also reanalyzed the 16S data via amplicon sequence variants (ASV) and found the 8th most abundant ASV to be *Nitrosopumilus*-like, while the next most abundant Thaumarchaeota ASV was ranked 754, confirming the dominance of one *Nitrosopumilus*-like species in the South Bay. However, the SSU in our dataset likely represents a different species than represented by the OTU, dominant ASV, or the two high-quality MAGs Rasmussen et. al. reported from these studies, based on an SSU alignment (Supplementary Figure 3). This suggests that there is still unexplored location or condition dependent dominance of specific *Nitrosopumilus* strains in the South Bay SFE.

Despite the existence of only one full length archaeal SSU, MMseqs2 classified contigs in 12 other archaeal phyla (Figure 2B). We note that the taxonomy output both by SILVA and MMseqs2 are not consistent, as the *Nitrosopumilus* contigs are classified under the Crenarchaeota and Thermoproteota, respectively, for the phylum, rather than Nitrososphaerota or Thaumarchaeota that is used by the International Committee on Systematics of Prokaryotes (Oren and Garrity 2021). Although there are only 14 *Nitrosopumilus* contigs (14 of the 21 Thermoproteota contigs in Figure 2B), the mean length of the contigs is twice that of the other archaeal contigs (24632 vs 12916bp). The longer contigs are consistent with the fact that the only full length archaeal SSU is from *Nitrosopumilus* and thus the assembly of this genome is more contiguous than the other archaeal genomes present.

We are not prepared to assert that the archaeal diversity is as great as the number of phyla listed above would indicate. We are using the GTDB taxonomy which is sometimes based on highly fragmented and incomplete bins and we consider some taxonomic collapse possible. However, coverage of the contigs involved is in the 5-10X range and thousands of bases in length and we believe there is a significant archaeal population at USGS Station 36. We consider it likely that much of the population is in the sediment rather than being planktonic and as part of future sequencing efforts, we will explore this further.

### Plasmids have a 3:1 ratio with archaeal and bacterial species

Based on the geNomad assignments, and removing the contigs that have SSUs, there are 1963 plasmids. Since we estimate that there are 450-500 archaeal and bacterial species (see next section for justification of this estimate), plasmids appear to be at a 3:1 ratio with the number of genomes. This ratio underscores the importance of ongoing efforts to link plasmids to chromosomes in metagenomes (Beaulaurier et al. 2018; Tourancheau et al. 2021). Of the plasmids that had MMseqs2 classification phylum level or below, three of the contigs were classified at the archaeal and 1591 were classified as bacterial, consistent with the estimated ratio of archaeal to bacterial species.

### Eukaryotic species, mitochondria, and chloroplasts

Using the eukaryotic specific database with the Kaiju classification of contigs, as well as manual curation of eukaryotic, mitochondrial, and chloroplast SSUs, allowed us to determine that we sequenced environmental DNA of large eukaryotes, such as mammals, polychaete worms, clams, mussels, sea anemone, zooplankton, ciliates, and diatoms (Supplementary Table 1, Supplementary Figure 4). The top 5 species based on contig counts classified by Kaiju were ciliate *Stylonychia lemnae* (896 contigs), diatom *Thalassiosira pseudonana* (524 contigs), green alga *Ostreococcus sp. ‘Lucimarinus’* (396 contigs), green alga *Picochlorum sp. BPE23* (273 contigs), and *Ostreococcus tauri* (272 contigs). Eleven species out of 256 comprised 82% of the eukaryotic contigs classified by Kaiju (Supplementary Table 3). The Kaiju database only had eukaryotic sequences from fungi and microeukaryotes, and did not report classifications from larger organisms.

We assembled some complete, circular mitochondria and chloroplast genomes and classified them based on the SSUs. In general, nearly all chloroplasts were initially classified as cyanobacteria (19/22) based on SILVA. One was classified as a *Marsupiomonas* chloroplast from manual curation. Only one mitochondrion was correctly classified by SILVA; the rest needed manual curation. We found that if we had only relied on the MMseqs2 classification of contigs, we would have misclassified the mitochondria and chloroplasts as bacteria. Most were classified as bacteria at the superkingdom level, but four of the mitochondria were classified to the species level (UBA4416 sp016787365, PWPS01 sp003554915, CAIVPM01 sp018335935) and two to the genus level (*Pelagibacter, UBA2645*). Similarly, there are some chloroplasts that are classified in the class Cyanobacteriia. Current theory holds that mitochondria arose from a bacteria becoming endosymbionts in an archaeal cell (Roger et al. 2017), so similarities of these genomes to bacterial ones aren’t unexpected. However, these results are a reminder to check if a MAG from a metagenome is a mitochondrion or chloroplast before labeling it as a new species.

### Estimation of number of prokaryotic genomes at USGS Station 36

Key questions in metagenomics concern what organisms are present in a sample and what the relative taxonomic abundances are. Not only do the answers to these questions provide an idea of the types of functions occurring in the sample, it also provides species abundances to analyze with environmental data. In order to estimate how many genomes we could aim to assemble, we used marker genes to estimate the number of species present and then used the coverage of the contigs as proxies for relative abundance.

Two common choices for marker genes are the small subunit (SSU) ribosomal RNA gene and 30S ribosomal protein S3 (rpsC) (Sharon et al. 2015). SSUs have the advantage that they are the basis for much of modern taxonomy (Woese and Fox 1977) prior to the advent of GTDB (Chaumeil et al. 2022) using full genomes or genome bins for taxonomic classification, and there are existing databases that can be used to place SSUs in the Tree of Life (Quast et al. 2013; Cole et al. 2014). At the same time, the disadvantages to using SSUs are that sometimes they have copy numbers greater than one, which distorts relative abundance measurements and, because they act as genomic repeats they fragment Illumina-based assemblies (see (Lui et al. 2021; Wick et al. 2017) for more detailed discussions about how repeats affect genome assembly). Using rpsC avoids the latter issue since the copy number is always one, but the lack of databases providing taxonomic information is problematic. Assemblies based on Nanopore reads can mitigate both issues with SSUs. Nanopore reads are long enough to span ribosomal operons and thus SSUs (and the rest of the ribosomal operon) do not cause assembly fragmentation (Koren and Phillippy 2015). Moreover, if the sequencing is deep enough, the contigs are often sufficiently long to contain both the genes for the SSU and for rpsC thus allowing rpsC genes to be assigned a taxonomy consistent with that of the SSUs. As we accumulate more nearly complete genomes, we intend to keep track of the correspondence between SSU and rpsC wherever possible.

To evaluate how much of the microbiome we were able to decompose into individual genomes, we used SSUs, rpsC, and contig taxonomy via MMSeqs2 (using GTDB as the database) to get a count of distinct organisms. First we evaluated the SSUs in the Nanopore assembly. We found a total of 561 bacterial and archaeal SSUs. Of those, 37 were reported as truncated by *cmsearch, i*.*e*., the covariance model couldn’t be fit at one or both ends. We manually curated the 37 and concluded that they were assembly artifacts or from extremely rare organisms and decided to exclude them from further consideration. This left 524 SSUs.

There are two confounding factors to using SSUs to estimate species abundances: (1) one genome can have multiple SSUs that have different sequences and (2) genomes from different species or strains can have identical SSUs (Jaspers and Overmann 2004). Accounting for contigs that contain multiple SSUs (29; Figure 3, Supplementary Table 2), the number of estimated species based on SSUs becomes 495. We found 17 contigs that have at least two different SSU sequences. Accounting for identical SSUs between different contigs, we found 49 groups of SSUs. We did not reduce the number of estimated species based on contigs having identical SSUs because it is not possible to tell if these contigs belong to the same genome or to different strains. For example, contig_37285 (772,808 bp) and contig_57621 (575,645 bp) both have 2 SSUs and all 4 of these SSUs are identical. However, an alignment of these two contigs using LAST (Kiełbasa et al. 2011) did not result in any significant alignment longer than 5,743 bp, which is likely the alignment of ribosomal operons. This suggests that these contigs either represent two pieces of the same genome or they are from different strains/species, despite having identical SSUs. For these reasons, some of the SSUs in Figure 3 are identical and our estimate of the number of species based on SSUs is still 495 but with a potential lower bound of 447.

**Figure 3:**
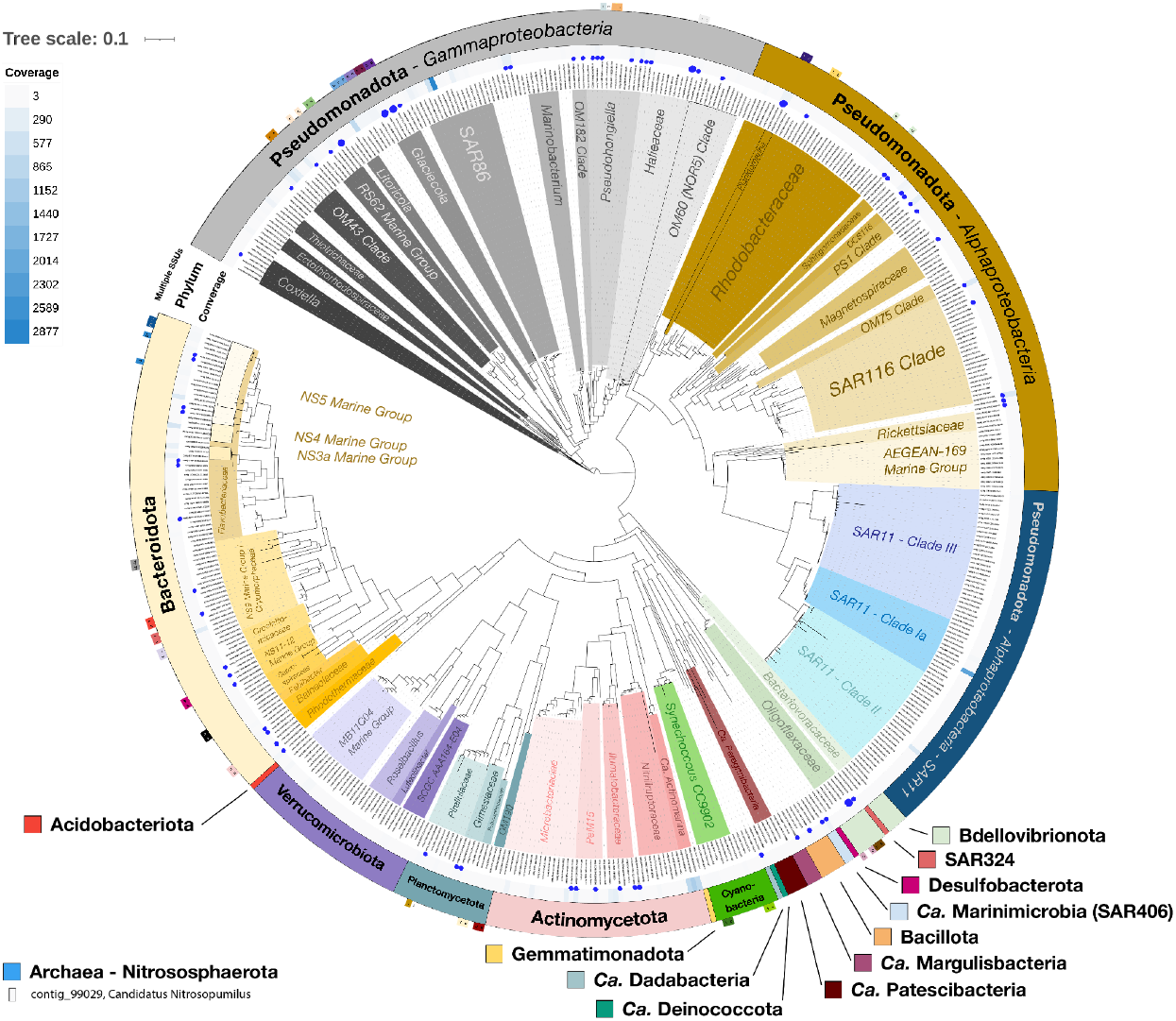
Taxonomy for Full-Length Bacterial and Archaeal SSUs. A tree of the bacterial SSUs is shown. There was only one full-length archaeal SSU, which is shown in the bottom left corner. The outer ring of the tree indicates if there is a pair of SSUs on the same contig (see Supplementary Table 2 for more detail), the next ring the phylum, and the third ring the coverage of the contig the SSU is on. The blue circles indicate that a high-quality bin exists with that SSU; larger circles are used for contigs that have more than one SSU.The inner taxonomy was added manually to highlight family, genus, or species level groups of interest. The coverage of the contig with the archaeal SSU is also indicated on the same scale as the bacterial tree (coverage of 8).

We next annotated all of the predicted protein-coding genes in the Nanopore assembly and extracted the rpsC genes. We found a total of 548 contigs with rpsC genes. However, the Nanopore assembly also contained genomes for 59 mitochondria and 23 chloroplasts, and these generally contain rpsC genes as well. Eighteen of the rpsC genes were on contigs known to have mitochondrial SSUs, so the upper bound on the number of prokaryotic species represented by rpsC genes is 530, and the lower bound is 466.

Finally, we used MMseqs2 taxonomy on all of our contigs with the GTDB database. We selected the contigs with species level taxonomic assignments, at least 10 labels that agreed with the contig assignment and coverage of at least 10 in order to get a conservative taxonomic assignment. This yielded an estimate of 489 as a lower bound for the number of distinct species. It must be understood that this taxonomic assignment amounts to a clustering of the longer contigs using the GTDB organisms as seeds. A set of contigs all being assigned to the same GTDB organism at the species level does not necessarily mean that it is the same species or even a strain; it may just be a statement that the contigs are closer to the GTDB organism than to anything else in the database and we infer that they are therefore likely to be close to each other as well.

Based on the three estimates, we conclude that the USGS Station 36 sample contained approximately 450-500 distinct bacterial organisms to which we can confidently assign taxonomy. Supplementary Table 2 lists the organisms in decreasing order of relative abundance, *i*.*e*., coverage of the contig. We also examined diversity based on the Illumina assembly and found a total of 467 bacterial SSUs. However, only 36 were not found to be truncated by *cmsearch*. It is worth noting that while the estimate based on the Illumina assembly leads to a consistent estimate of the number of distinct species, it doesn’t generally allow assignment to lower taxonomic ranks. Most of the fragments were significantly shorter than 1,000 bp and only the extreme sensitivity of *cmsearch* allowed them to be seen (Nawrocki and Eddy 2013).

We generated a phylogenetic tree based on the bacterial SSUs to study the phylogeny, and included coverage of the contig the SSU was on as a proxy for abundance (Figure 3). Six species comprised 30% of the relative abundance of bacteria and archaea: a SAR86 species (10.3%), a SAR11 Clade Ia species (7.8%), three *Actinomarina* species (3.9%, 3%, 2.9%), and a *Litoricola* species (2.8%). Notably, all of these species except for the *Actinomarina* had closely related relatives that were not highly abundant. These results suggest that in order to study the population dynamics of the estuary microbiome, we will need to look at the individual genome level. Although we were able to tell which species were the most abundant, obtaining high-quality genome bins is necessary to help understand why some may be more successful than others.

### Binning of Prokaryotic Genomes and Bin Refinement

Ideally, we would like to extract the genome of each of the ∼500 bacteria and archaea we believe to be present at Station 36 as one - generally circular - chromosome per contig. A total of 6 genomes were circularized immediately by the assembler. We added a few by inspection. But the bulk of the genomes are present in the assembly as a collection of fragmented contigs. This is what binning software is designed to deal with. Unfortunately, most binning software is designed to be used with Illumina based assemblies and may not work well with Nanopore assemblies due to the higher error rates. We looked into several recent efforts to produce long read binning software and we eventually settled on GraphMB (Lamurias et al. 2022). To obtain high quality genomes, we did initial binning with GraphMB, and then did bin refinement by a novel method (Figure 1C).

We used GraphMB with default parameters on the Nanopore assembly, and then refined the list of bins to find candidates for refinement. GraphMB produced a total of 1,310 bins. We used CheckM2 (Chklovski et al. 2022) to evaluate the bins to see which ones fit our criteria for nearly complete genomes. Of the total, 128 bins met the requirement that completion was greater than 90%. Of those that did, 59 also met the requirement that the contamination was less than 5%. The remaining 69 had contamination greater than 5%. All of the genomes reported to be circular by Flye were included in the 128.

Next, we used a novel bin refinement method on these 128 bins. For short read metagenomes, we previously published an algorithm called Jorg for improving bins to the point where they consisted of a single circular chromosome (Lui et al. 2021). We adapted the workflow used there to improve the bins. For each of the bins meeting the completion requirement, we used minimap2 (Li 2018) with default parameters to map the full set of long reads to the contigs in the bin. We then used the reads thus obtained as input to Canu (Koren et al. 2017). Canu is an overlap consensus assembler and generally produces fewer misassemblies than Flye, but because of the computational requirements is too difficult to run on the entire set of nanopore reads. We ran Canu with default parameters to produce a set of contigs which we then manually curated based on GC, coverage and CheckM2 results to obtain a new bin. We then iterated the process using the new bin instead of the old one.

We terminated each iteration when there was no further clear improvement in either completion, contamination or number and length of contigs. For 68 of the original bins, we were able to improve them to where they met our definition of nearly complete genomes, *i*.*e*., >90% complete, <5% contamination, no more than 5 contigs, a full ribosomal RNA gene complement, all necessary tRNAs and no contigs shorter than 100 Kbases (Supplementary Table 4). Anecdotally, we would like to point out that of all 128 bins we worked on, only 1 failed due to the requirement of no more than 5 contigs and no contig shorter than 100 Kbases implying that in practice, these are not particularly onerous requirements. Almost all of the failures were due to inadequate coverage.

This bin refinement method allows the merging of contigs and the separation of highly similar strains. The example of contig_37285 and contig_57621 in the previous section, where both contigs have identical SSUs, demonstrated that these two contigs belong to the same genome. In other cases, we found that two genomes would separate that have identical SSUs, but whole genome alignments indicate that they should be separate strains. We should also point out that the main determinant for success appears to be coverage. If coverage is adequate - at least ∼20X - our bin improvement method works extremely well. The exception to that rule is SAR11. SAR11 organisms are known to be very difficult to bin (López-Pérez et al. 2020) and we expect to need significantly more sequencing and possibly isolation efforts to complete their genomes.

The methodology used for generating bins here is extremely flexible. Using it with contigs where coverage is ∼40X or better, improvement is generally swift. A significant number of bins are high quality drafts at the start and only need a reduction in the number of contigs and a restriction on the length of the contigs. A number of the nearly complete genomes we have are present as a single contig. We have also experimented with using single or a few long contigs we believed to belong together as starting material and then iterated to extend the contigs. In some cases, the process can be accelerated by looking for significant overlaps with other contigs in the assembly.

The biggest issue with our methodology is the amount of manpower it requires. We estimate that producing a nearly complete genome like this takes 5-10 man hours per genome. However, the quality of the final result is worth it to us. Our goal is to completely decompose the metagenomes in the SFE and to obtain complete genomes for as many of them as possible. As we move forwards and sequence more, we expect the work we do now will allow us to reduce the time investment needed. We have decided to only report on only nearly complete genomes because of the issues that lower quality bins can cause in downstream analyses.

### Viruses Compose ∼43% of the Station 36 contigs

Marine viruses have a huge influence on the population dynamics of marine microbial populations, as well as element cycling in the marine environment (Breitbart et al. 2018). As mentioned in a previous section, viruses comprise a large part of the total contigs in our sample (Figure 1A). We cannot account for RNA viruses in this analysis, so the ratio of viruses to the other organisms in the sample is likely even higher. Using geNomad, 45,790 contigs were annotated as viruses. Two of these had SSUs, so we reduced this number to 45,788. Given that the assembly had 107,977 contigs in total, viruses account for 42.4% of the contigs. Considering bases rather than contigs produces a very similar number. We did not expect such a high proportion of viruses and we attribute it to using a 0.1 μm filter and letting it come very close to clogging.

To estimate the number of viral populations, we used the heuristic of ≥95% ANI as determined by an analysis of the Global Oceans Viromes 2.0 (GOV 2.0) dataset (Gregory et al. 2019). There were 18 sequences that were considered duplicates at ≥95% ANI and 611 sequences that were contained in other sequences, so we estimate the total number of viral populations to be 45,169 in our dataset. Of these, 3,112 were found to be circular by the Flye assembler. The median length of the viral population sequences is 11,624 bp, and the mean is 23,243bp (Figure 4). Despite the modest average length of the viral population sequences, there are 88 longer than 300 kbp, 3 of which are greater than 1Mbp, which puts them into the genome size of giant viruses (Schulz et al. 2022). Of these, 20 are in the Order *Caudoviricetes* and 14 are circular (Supplementary Figure 5). The rest are in the Phylum *Bamfordvirae*, where all except one are classified in the Order *Megaviricetes*, and 8 are circular. The two longest circular viruses are 601,220 and 1,006,597 bp and their lowest taxonomic classification is *Megaviricetes*.

**Figure 4:**
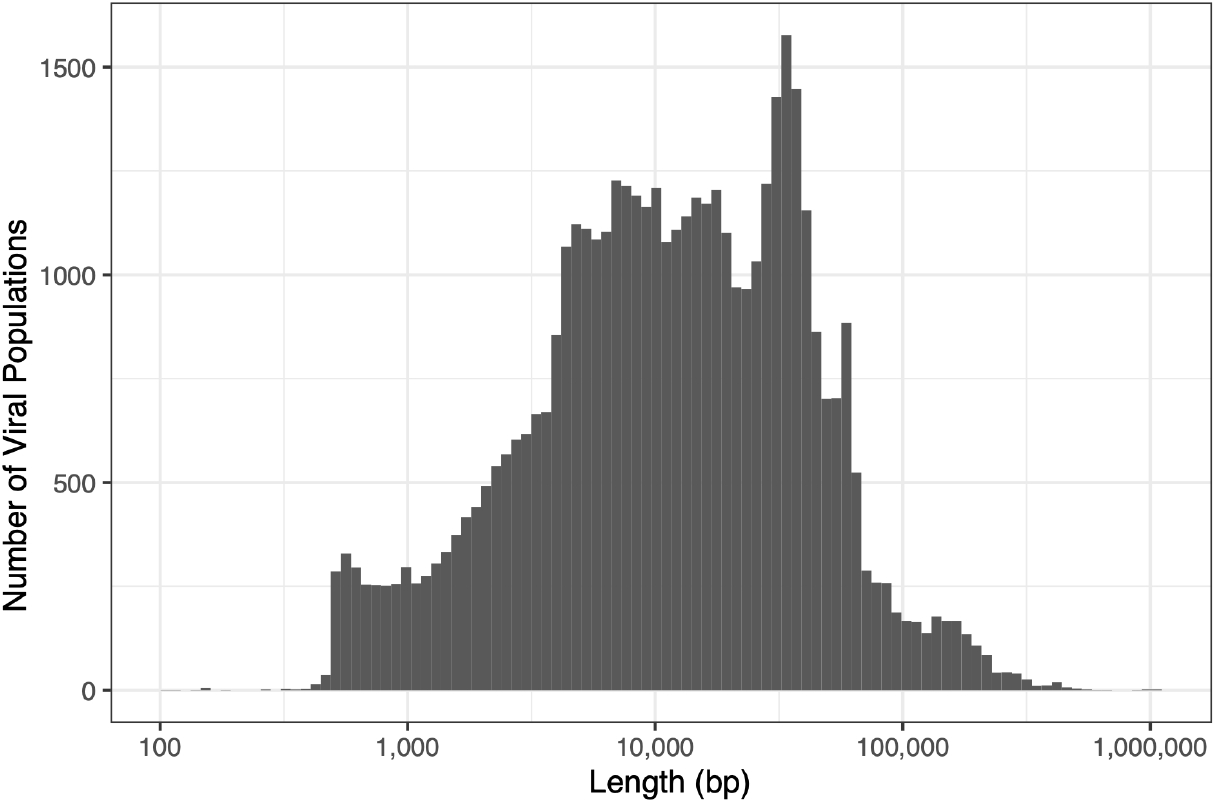
Histogram of Viral Population contig length in the Station 36 metagenome.

There were three viral Realms represented in the dataset (Supplementary Figure 6), although 5281 could not be classified at the Realm level and two populations were classified in the Class *Naldaviricetes*, which does not have a higher taxonomic classification assigned to it. The Realm *Duplodnaviria* composed 80% of the viral populations, 99.9% of which was classified as *Caudoviricetes* (Supplementary Figure 6B). Only 2 viral populations were classified under the Realm *Monodnaviria* (Supplementary Figure 6C). In the Realm *Varidnaviria*, 3787 populations were present in the sample, and 58% of this Realm was classified under the Family *Phycodnaviridae* (Supplementary Figure 6D).

The *Phycodnaviridae* contigs comprise approximately 4.8% of the total viral populations, and members of this Family are the most abundant viral populations in this sample (Figure 5). Members of this Family are known to infect marine picoeukaryotic and microplankton species, such as *Ostreococcus*, and diatoms (Schulz et al. 2022), so it is not unexpected that these would be highly abundant in our sample, and have been found to be abundant in other estuarine viromes (Sun et al. 2021). Twenty of the 2,195 *Phycodnaviridae* populations have coverage >1,000 and range in length from ∼31-236 kbp (Supplementary Figure 7). The dominance of 10% of the *Phycodnaviridae* populations in terms of abundance emphasizes the importance of finding a way to determine the top abundant viral strains in a sample.

**Figure 5:**
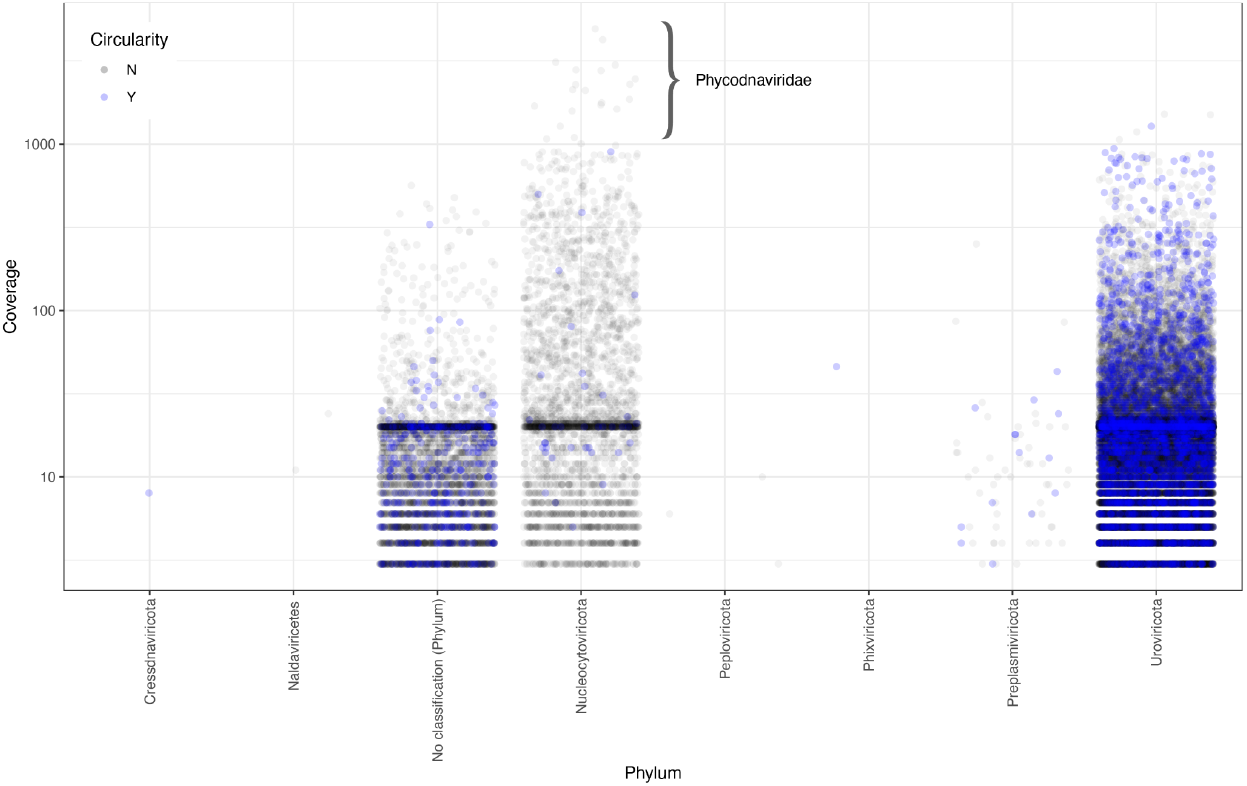
Plot of coverage of the viral populations by phylum. Circular viral populations are colored in blue. In the *Nucleocytoviricota*, all of the viral populations with coverage >1000 belong to the *Phycodnaviridae* Family except one.

Determining the hosts of viruses is still difficult, but is extremely important for modeling the marine microbiome population dynamics (Breitbart et al. 2018). We examined the intersection of the MMSeqs2 taxonomy with the geNomad predictions more closely, especially since 38,109 contigs of the 46,478 predicted to be viral by geNomad were classified as either archaea or bacteria by MMseqs2. We found that the top 5 longest of viral contigs classified as archaea by MMseqs2 were all circular and between 289,857 to 307,949 bp in length. Manual analysis of these contigs indicated that these were indeed archaea phage, as the presence of viral genes were spread through the entirety of the contigs. Specifically the MMSeqs2 classification for all 5 is “d_Archaea; p_Nanoarchaeota; c_Nanoarchaeia; o_Pacearchaeales; f_GW2011-AR1”, which suggests that these are a type of Nanoarchaeota phage. It is possible that we can use some of the MMSeqs2 classification to help determine hosts of some of these viruses, but will require further analysis that is beyond the scope of this study. Plotting coverage of viruses by taxonomy indicates that there may be phage strain specific dominance (Supplementary Figure 8), based on this hypothesis.

## Conclusion

In this study, we have expended significant effort on determining exactly who is there down to the species/strain level. One of our major goals is to fully resolve the metagenomes into their constituent genomes and to extract complete genomes for as many of the organisms as possible. In general, 20x coverage of a genome is considered necessary to ensure a high-quality assembly, at least based on analysis of Illumina metagenomes (Luo et al. 2012). Thus, to fully decompose this metagenome, we need to have at least six times more sequencing depth (3x was the lowest coverage of any full length SSU), so approximately 1Tbase. We are planning to sequence deeper in the future.

This is just one sample. Our goal is to sequence the bacterial and archaeal microbiome of the San Francisco Estuary to obtain complete genomes for all of the constituent organisms. This will require a substantial sequencing effort. Our initial tranche is 5 Tbases of Nanopore long read sequencing. We expect to (1) catalog the microbial diversity in the SFE, (2) generate fully complete genomes of as many of the microbial species in the SFE as possible, and (3) catalog mobile genetic elements in the SFE, such as viruses and plasmids. As this project unfolds, we expect to have enough data to produce a database that assigns taxonomy to rpsC genes based on the taxonomy of their SSUs. This is just one example of what we expect to be able to do with the volume of data. Because we sequenced deeply with Nanopore reads, the assembly also had contigs of sufficient length to assign taxonomy to a large percentage of them using tools such as MMseqs2 (Mirdita et al. 2021). There have been multiple calls for long term monitoring and study of marine microbiomes (Cavicchioli et al. 2019; Brennan and Logares 2023), and this study is a first step to monitoring changes at the genome level.

## Supporting information

Classification

SSU Data

Kaiju Results

## Acknowledgements

We would like to thank USGS for collecting samples for us, especially Erica Nejad and the crew of USGS R/V David H. Peterson. We cannot overemphasize the value of the support we have received for this project from USGS. We also would like to thank the support that we received from the Joint Genome Institute for PromethION sequencing and library quality analysis. Specifically we would like to thank Hope Hundley, Rob Egan, Len Pennacchio, and Ronan O’Malley.

This work was supported by the Laboratory Directed Research and Development Program of Lawrence Berkeley National Laboratory under U.S. Department of Energy (DOE) Contract No. DE-AC02-05CH11231.

Part of this work (proposal:https://doi.org/10.46936/10.25585/60008607) was conducted at the U.S. DOE Joint Genome Institute (https://ror.org/04xm1d337), a DOE Office of Science User Facility supported by the Office of Science of the U.S. Department of Energy operated under Contract No. DE-AC02-05CH11231.

