## Supplementary Data for "Decomposing a San Francisco Estuary microbiome using long read metagenomics reveals species and species- and strain-level dominance from picoeukaryotes to viruses"

### Supplementary Tables and Figures

**Supplementary Table 1: SSU Taxonomic Classification for Bacteria, Archaea, Eukarya, Mitochondria, and Chloroplasts.** Coverage values refer to the average coverage of the contig that the SSU is on. SILVA and manual classification by NCBI BLAST search included.

**Supplementary Table 2: Bacterial and Archaeal SSU Classification by MMSeqs2 using GTDB as the Database.** SSUs that are on the same contig are indicated by color and letter that match annotations in Figure 3.

**Supplementary Table 3: Kaiju Classification of the Eukaryotic Contigs.**

**Supplementary Table 4: Bin Quality and Taxonomic Classification.** Results for CheckM2 completeness and contamination and taxonomic classification by GTDB-tk are included.

**Supplementary Figure 1: Electropherogram of Femto Pulse separation of size-selected DNA from USGS Station 36.**

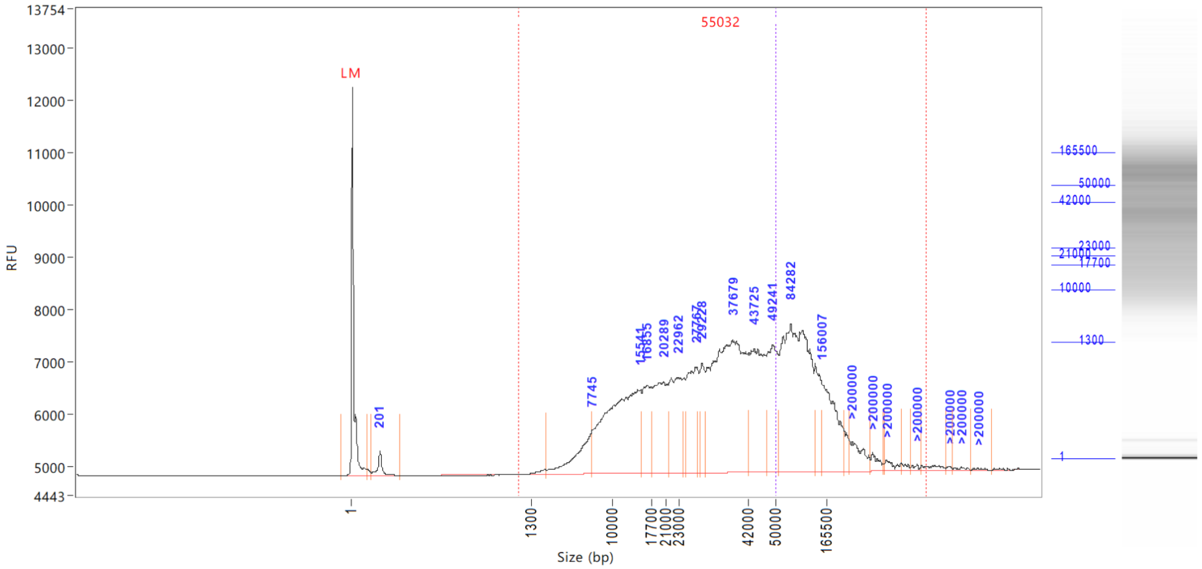


**Supplementary Figure 2: Femto Pulse of Nanopore library made with SQK-LSK112 kit.**

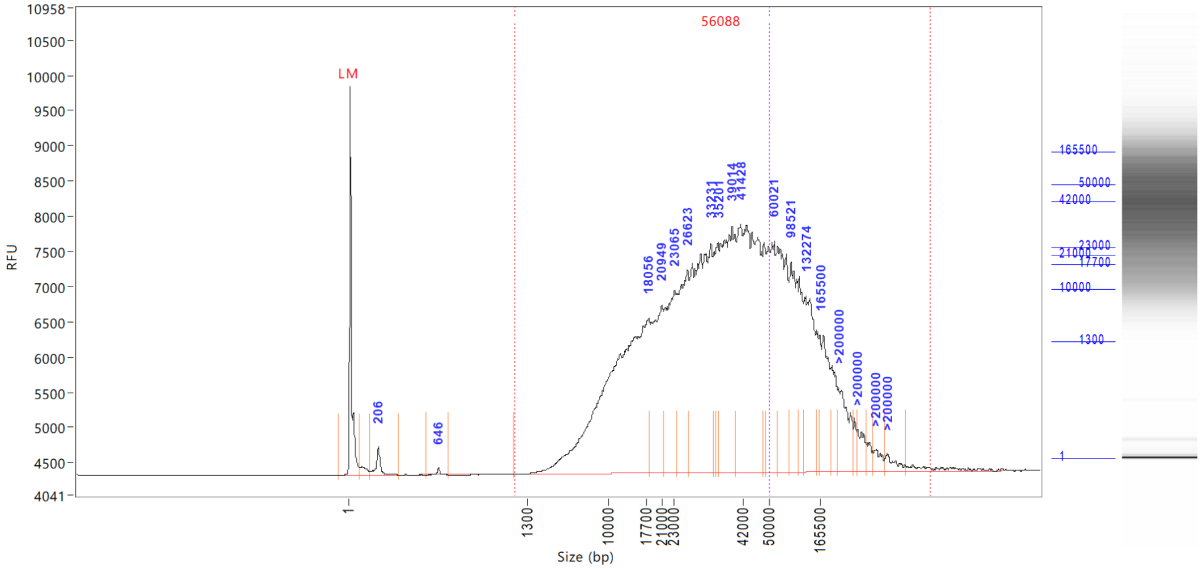

**Supplementary Figure 3: Alignment of *Nitrosopumilus* SSU with SSUs from Rasmussen et al. 2022.** Rasmussen et al. 2021 Microbial Ecology reported on dominant archaeal OTU (Thaumarchaeota) from the South Bay of the San Francisco Estuary. They also subsequently reanalyzed the 16S data via ASVs in a paper focused on two ammonia-oxidizing archaea MAGs. The ASV analysis indicated that of the Thaumarchaeota ASVs, the most abundant ranked 8th of all ASVs (ASV8), while the next most abundant Thaumarchaeota ASV ranked 754. This suggests that this ASV is similar to the dominant OTU. We wanted to see if the SSU in our dataset matched either the ASV or OTU. Since the OTU and ASV sequences were not published from Rasmussen et al, we aligned our SSU with SSUs from two ammonia-oxidizing archaea MAGs that Rasmussen et al assembled from USGS Station 3 and 27 water samples. The Station 3 MAG had one full length SSU, while the Station 27 MAG had one full length SSU and one 568 bp fragment. Rasmussen et al. report that the Station 27 MAG exactly matches ASV8 from their 16S study, which we assume is to the full length SSU, as the fragment does not cover the entirety of the V4-V5 region. The SSU from our dataset had only ~86% identity to both full length SSUs and ~91% identity to the fragment, so it is likely a different genus or species (Yarza et al. 2014).

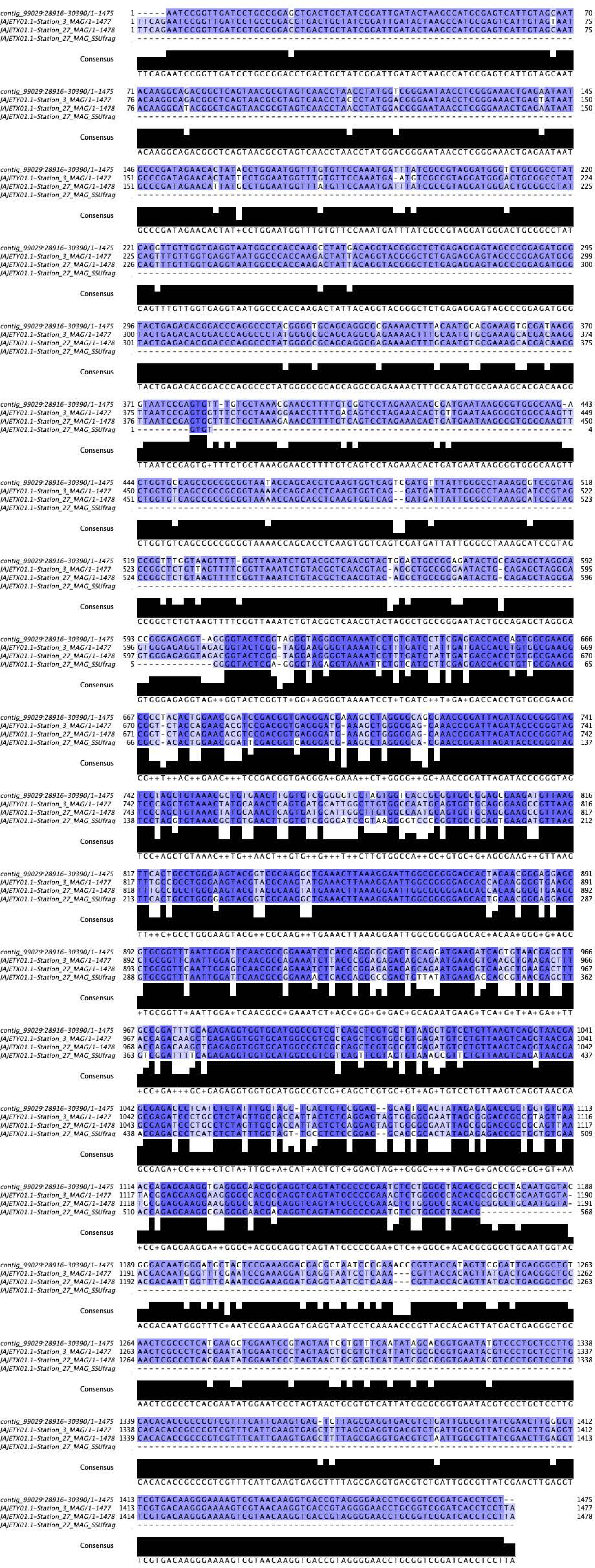
contig_99029:28916-30390/1-1475
JAJETY01.1-Station_3_MAG/1-1477
JAJETX01.1-Station_27_MAG/1-1478
JAJETX01.1-Station_27_MAG_SSUfrag
Consensus

**Supplementary Figure 4: Eukaryotic phyla of contigs classified by Kaiju and number of contigs detected for each species.** Taxonomic class is listed on the x-axis and species with >100 contigs are labeled with the genus.

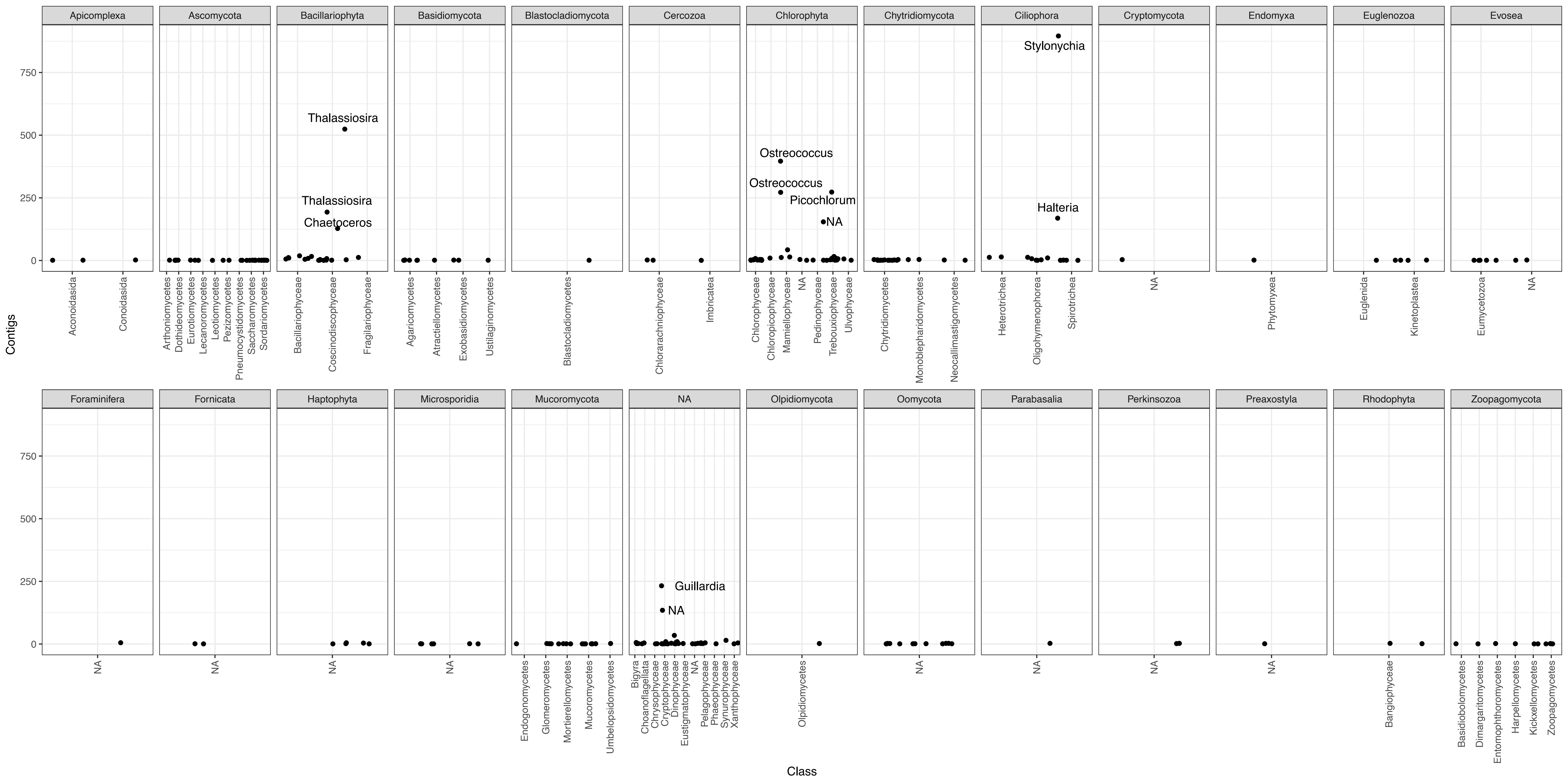

Contigs
Class
0
250
500
750
Apicomplexa
Aconoidasida
Conoidasida
Ascomycota
Arthoniomycetes
Dothideomycetes
Eurotiomycetes
Lecanoromycetes
Leotiomycetes
Pezizomycetes
Pneumocystidomycetes
Saccharomycetes
Sordariomycetes
Bacillariophyta
Thalassiosira
Thalassiosira
Chaetoceros
Bacillariophyceae
Coscinodiscophyceae
Fragilariophyceae
Basidiomycota
Agaricomycetes
Atractiellomycetes
Exobasidiomycetes
Ustilaginomycetes
Blastocladiomycota
Blastocladiomycetes
Cercozoa
Chlorarachniophyceae
Imbricatea
Chlorophyta
Ostreococcus
Ostreococcus
Picochlorum
NA
Chlorophyceae
Chloropicophyceae
Mamiellophyceae
NA
Pedinophyceae
Trebouxiophyceae
Ulvophyceae
Chytridiomycota
Chytridiomycetes
Monoblepharidomycetes
Neocallimastigomycetes
Ciliophora
Stylonychia
Halteria
Heterotrichea
Oligohymenophorea
Spirotrichea
Cryptomycota
NA
Endomyxa
Phytomyxea
Euglenozoa
Euglenida
Kinetoplastea
Evosea
Eumycetozoa
NA
Foraminifera
NA
Fornicata
NA
Haptophyta
NA
Microsporidia
NA
Mucoromycota
Endogonomycetes
Glomeromycetes
Mortierellomycetes
Mucoromycetes
Umbelopsidomycetes
NA
Guillardia
NA
Bigyra
Choanoflagellata
Chrysophyceae
Cryptophyceae
Dinophyceae
Eustigmatophyceae
NA
Pelagophyceae
Phaeophyceae
Synurophyceae
Xanthophyceae
Olpidiomycota
Olpidiomycetes
Oomycota
NA
Parabasalia
NA
Perkinsozoa
NA
Preaxostyla
NA
Rhodophyta
Bangiophyceae
Zoopagomycota
Basidiobolomycetes
Dimargaritomycetes
Entomophthoromycetes
Harpellomycetes
Kickxellomycetes
Zoopagomycetes

**Supplementary Figure 5: Coverage vs Length of Giant Viruses in Station 36 metagenome**

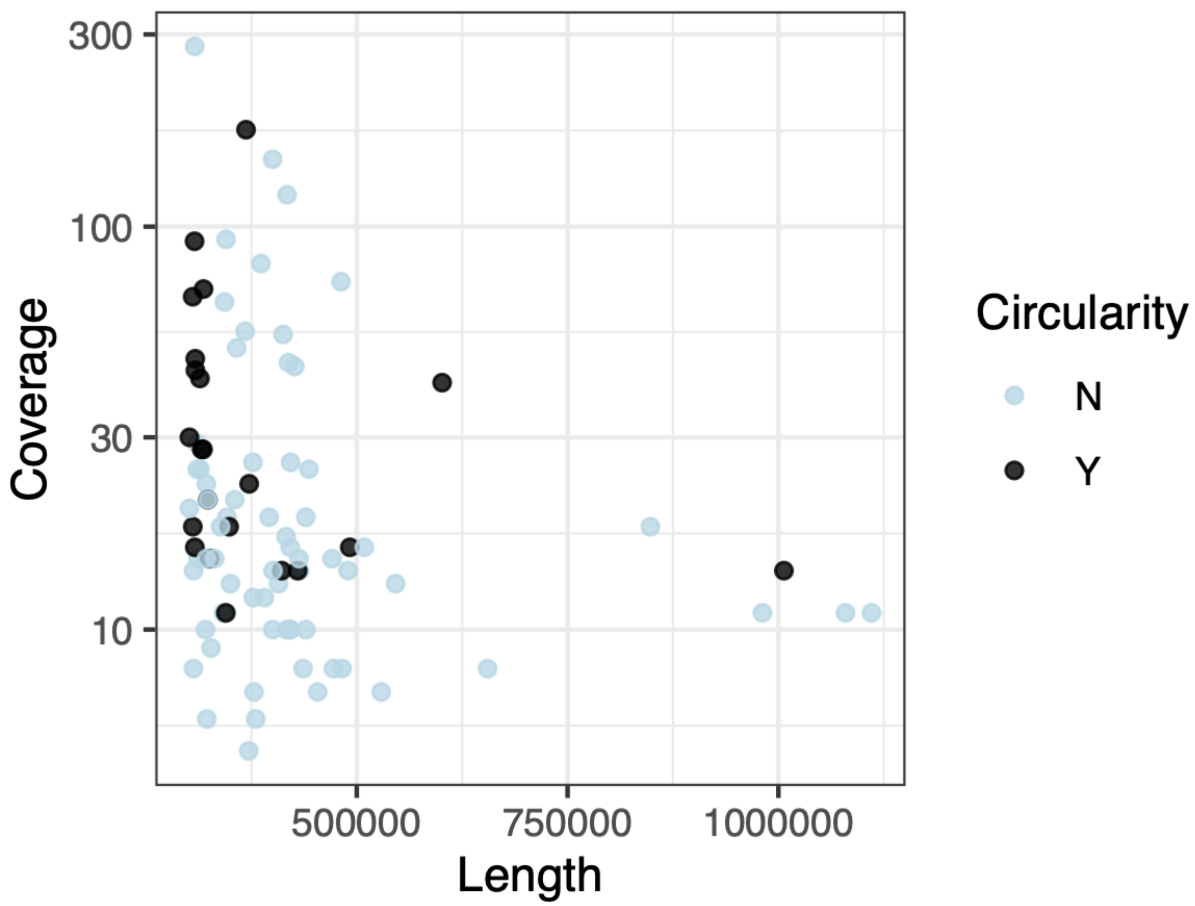

**Supplementary Figure 6: Viral Taxonomy of the Station 36 metagenome.** (A) Viral Realms detected. (B) Breakdown of Realm *Duplodnaviria.* (C) Breakdown of Realm *Monodnaviria*. (D) Breakdown of Realm *Varidnaviria*.

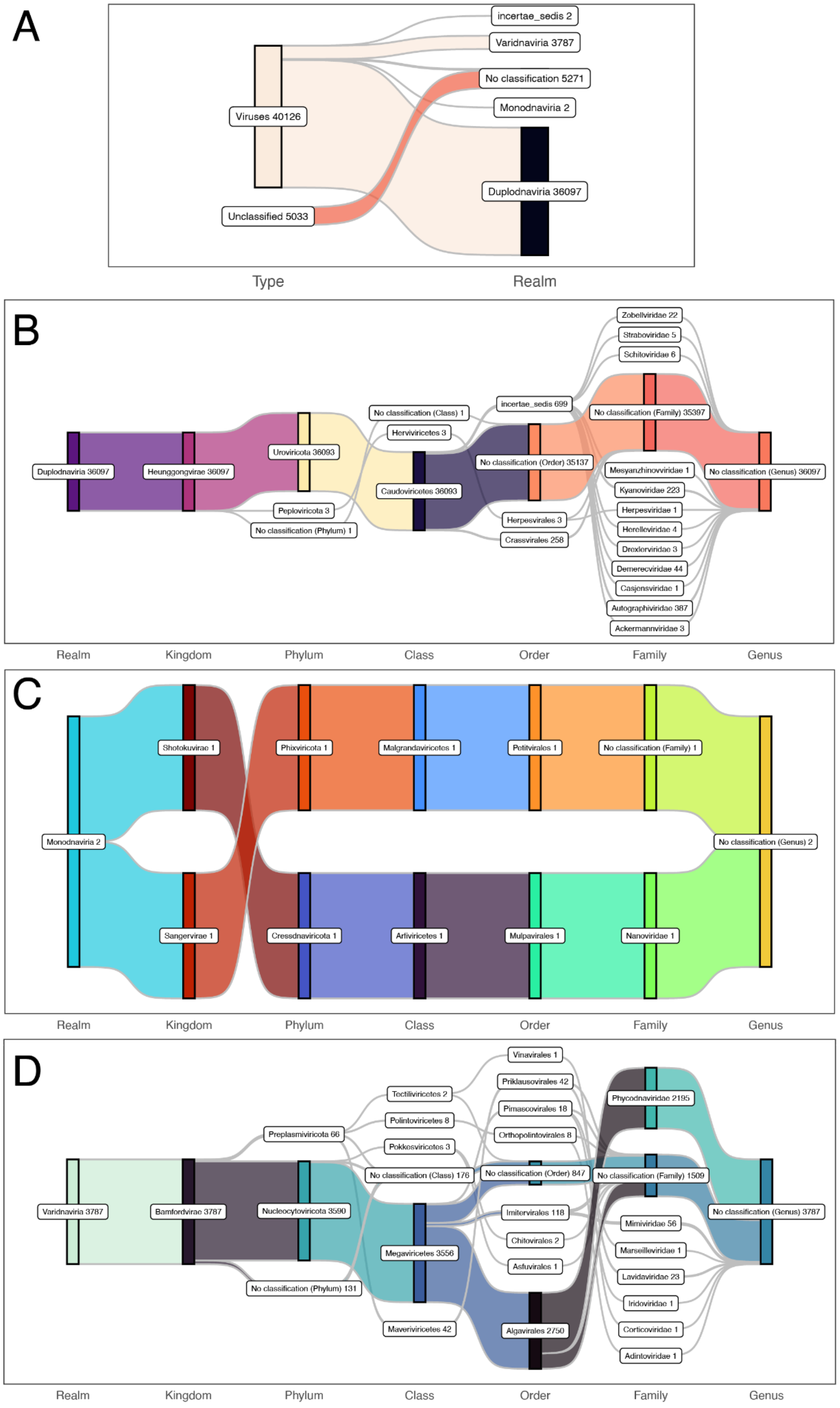

**Supplementary Figure 7: *Phycodnaviridae* populations plotted by coverage and length (bp).**

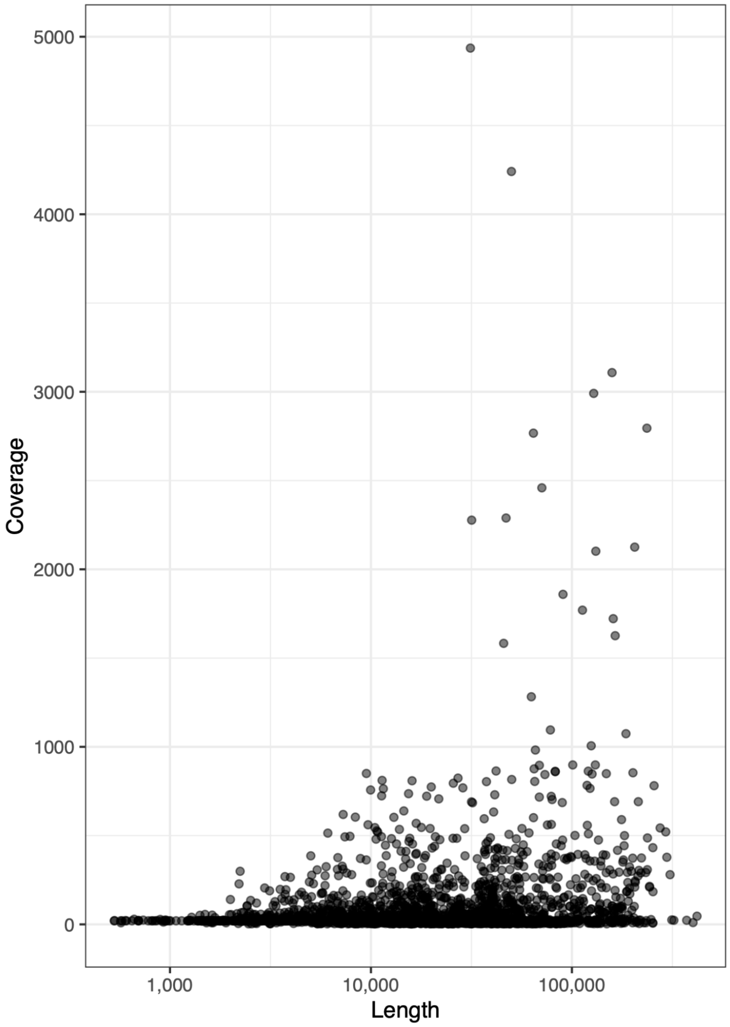

**Supplementary Figure 8: *Proteobacteria* phage populations plotted by coverage and taxonomic family classification.**

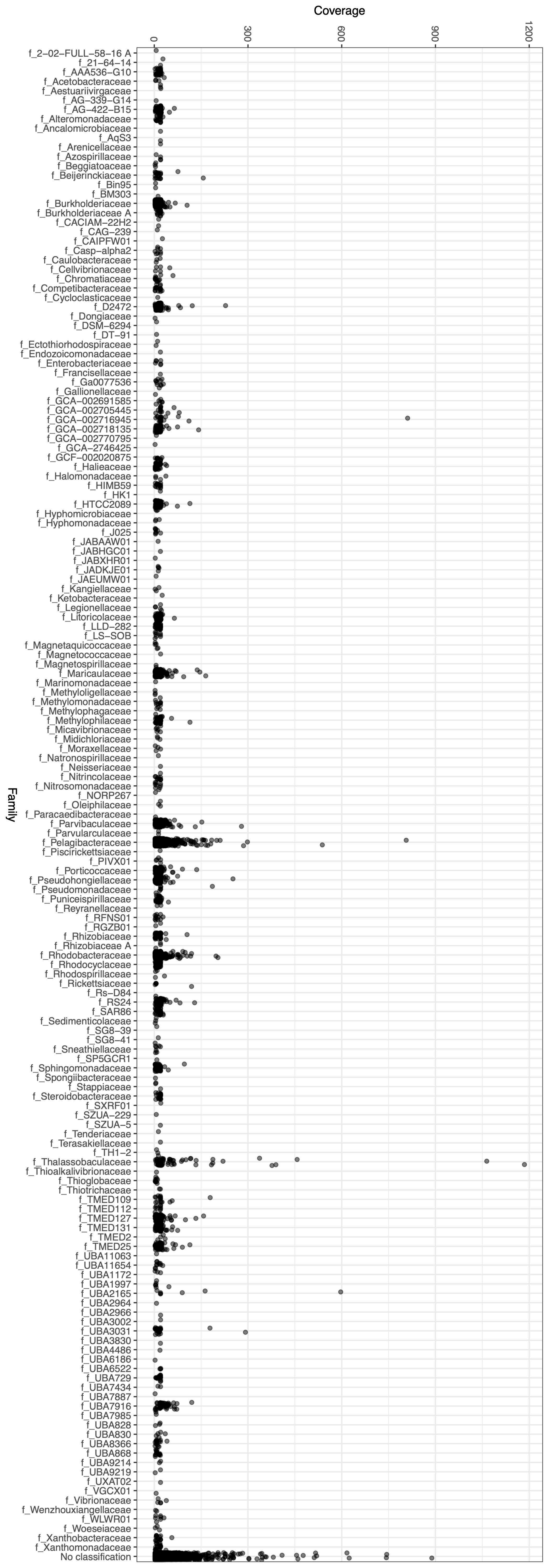

Coverage
0
300
600
900
1200
Family
f_2-02-FULL-58-16 A
f_21-64-14
f_AAA536-G10
f_Acetobacteraceae
f_Aestuariivirgaceae
f_AG-339-G14
f_AG-422-B15
f_Alteromonadaceae
f_Ancalomicrobiaceae
f_AqS3
f_Arenicellaceae
f_Azospirillaceae
f_Beggiatoaceae
f_Beijerinckiaceae
f_Bin95
f_BM303
f_Burkholderiaceae
f_Burkholderiaceae A
f_CACIAM-22H2
f_CAG-239
f_CAIPFW01
f_Casp-alpha2
f_Caulobacteraceae
f_Cellvibrionaceae
f_Chromatiaceae
f_Competibacteraceae
f_Cycloclasticaceae
f_D2472
f_Dongiaceae
f_DSM-6294
f_DT-91
f_Ectothiorhodospiraceae
f_Endozoicomonadaceae
f_Enterobacteriaceae
f_Francisellaceae
f_Ga0077536
f_Gallionellaceae
f_GCA-002691585
f_GCA-002705445
f_GCA-002716945
f_GCA-002718135
f_GCA-002770795
f_GCA-2746425
f_GCF-002020875
f_Halieaceae
f_Halomonadaceae
f_HIMB59
f_HK1
f_HTCC2089
f_Hyphomicrobiaceae
f_Hyphomonadaceae
f_J025
f_JABAAW01
f_JABHGC01
f_JABXHR01
f_JADKJE01
f_JAEUMW01
f_Kangiellaceae
f_Ketobacteraceae
f_Legionellaceae
f_Litoricolaceae
f_LLD-282
f_LS-SOB
f_Magnetaquicoccaceae
f_Magnetococcaceae
f_Magnetospirillaceae
f_Maricaulaceae
f_Marinomonadaceae
f_Methyloligellaceae
f_Methylomonadaceae
f_Methylophagaceae
f_Methylophilaceae
f_Micavibrionaceae
f_Midichloriaceae
f_Moraxellaceae
f_Natronospirillaceae
f_Neisseriaceae
f_Nitrincolaceae
f_Nitrosomonadaceae
f_NORP267
f_Oleiphilaceae
f_Paracaedibacteraceae
f_Parvibaculaceae
f_Parvularculaceae
f_Pelagibacteraceae
f_Piscirickettsiaceae
f_PIVX01
f_Porticoccaceae
f_Pseudohongiellaceae
f_Pseudomonadaceae
f_Puniceispirillaceae
f_Reyranellaceae
f_RFNS01
f_RGZB01
f_Rhizobiaceae
f_Rhizobiaceae A
f_Rhodobacteraceae
f_Rhodocyclaceae
f_Rhodospirillaceae
f_Rickettsiaceae
f_Rs-D84
f_RS24
f_SAR86
f_Sedimenticolaceae
f_SG8-39
f_SG8-41
f_Sneathiellaceae
f_SP5GCR1
f_Sphingomonadaceae
f_Spongiibacteraceae
f_Stappiaceae
f_Steroidobacteraceae
f_SXRF01
f_SZUA-229
f_SZUA-5
f_Tenderiaceae
f_Terasakiellaceae
f_TH1-2
f_Thalassobaculaceae
f_Thioalkalivibrionaceae
f_Thioglobaceae
f_Thiotrichaceae
f_TMED109
f_TMED112
f_TMED127
f_TMED131
f_TMED2
f_TMED25
f_UBA11063
f_UBA11654
f_UBA1172
f_UBA1997
f_UBA2165
f_UBA2964
f_UBA2966
f_UBA3002
f_UBA3031
f_UBA3830
f_UBA4486
f_UBA6186
f_UBA6522
f_UBA729
f_UBA7434
f_UBA7887
f_UBA7916
f_UBA7985
f_UBA828
f_UBA830
f_UBA8366
f_UBA868
f_UBA9214
f_UBA9219
f_UXAT02
f_VGCX01
f_Vibrionaceae
f_Wenzhouxiangellaceae
f_WLWR01
f_Woeseiaceae
f_Xanthobacteraceae
f_Xanthomonadaceae
No classification

### Supplementary Methods

#### Taxonomic classification of contigs at the Superkingdom level

As described in the main text, to classify the taxonomy of contigs we used geNomad, mmSeqs2, Kaiju, and manual curation based on SSUs and BLAST hits. In total, the assembly consisted of 107,977 contigs.

1. geNomad results:
    - 1963 Plasmids
    - 46476 Viruses
    - Notes: Viral taxonomy is discussed in the main text
2. MMseqs2 results (using GTDB as the database):
    - 85445 Bacteria
    - 1646 Archaea
    - 6493 Root
    - 1698 No rank
    - 12695 Unclassified by mmseqs2
3. Kaiju results (using nr_euk as the database):
    - 44322 Bacteria
    - 751 Archaea
    - 5822 Eukaryota
    - 23499 Viruses
    - 8176 Cannot be assigned to a non-viral species
    - 25407 Unclassified
4. Manual curation of SSU containing contigs:
    - 493 Bacteria
    - 1 Archaea
    - 69 Eukaryota
    - 23 Chloroplasts
    - 59 Mitochondria
    - Notes: This list only contains contigs with full length SSUs to ensure accuracy. These numbers also take into account contigs that have more than one SSU (see Supplementary Table X).

We had a number of assumptions about the classifiers that influenced how we interpreted the results:

1. **geNomad was the best at classifying contigs as viral or plasmid given the careful curation of plasmid and viral sequences going into the training data.** This assumption is also partially because first, we did not focus on using Kaiju to classify viruses and did not use their viral specific database when running Kaiju on our contigs, and second, Kaiju accuracy was not benchmarked on classification of viral sequences.

Most Kaiju benchmarks were on sensitivity and accuracy of prokaryotic phylogeny (see Menzel et al 2016. Nat. Communications). We also assume that some of the MMseqs2 contigs were classified as bacterial or archaeal because often viruses pick up genes from their hosts.

2. **MMseqs2 is the best at classifying contigs as bacterial or archaeal, except if the contigs is viral, and Kaiju will be better at classifying contigs as eukaryotic, because of the databases used**. Given that we used the GTDB database and MMseqs2 algorithm relies on finding specific proteins in a contig and determining if there is a close match in the database, we consider this algorithm more accurate at prokaryotic classification than Kaiju, at least if a close genome exists in the GTDB database and the contig is long enough to contain enough identifying genes. For bacterial and archaeal genomes, the GTDB database is more highly curated than the Kaiju database we used (contains sequences from NCBI BLAST nr database of bacteria and archaea), and thus is likely more accurate for those taxonomies. Since we used the Kaiju database that includes eukaryotic sequences, it will automatically be more accurate at classifying contigs as eukaryotic than the MMseqs2 results.
3. **In all cases, we assume that sequences that were only classified at the superkingdom level (*i.e.,* a sequence classified as Bacteria with no phylum or lower taxonomic rank assigned) are ambiguous.** Taxonomy of sequences which do not have a close genome in the database (such as the genus or family level), will depend on the last common ancestor (LCA) algorithm used, and these are different between Kaiju and MMseqs2. We do not take a stance on which of these algorithms are more accurate.

First, we started with the contigs that we manually curated based on their SSUs, along with SILVA and BLAST taxonomy assignments. Since we manually curated these, we consider these accurate. We consider chloroplasts and mitochondria as eukaryotic.

- 493 Bacteria
- 1 Archaea
- 151 Eukaryota
- 107332 Unclassified

Next we added in the geNomad results for the viruses and plasmids. Eight contigs that were assigned as a plasmid and one that was assigned as a virus had SSUs, so these contigs were excluded from the counts.

We started with this because many of the sequences classified as viral by geNomad were classified as archaeal or bacterial by MMseqs2, likely because some of these sequences also have some archaeal and bacterial genes. It is possible that some of the geNomad classification is not accurate, but we do not have an easy way to evaluate the accuracy, short of searching every contig for hallmark viral genes and manual curation. Of the 46475 contigs that were classified as viral by geNomad, 3697 were unclassified, 18734 were classified at the superkingdom level, 4231 as root, and 441 had no rank, so 27103 (58%) of these sequences we would already consider ambiguous already by MMseqs2 assignments. So this number would be a more conservative accounting for the number of viruses, and we keep this in mind going forwards with the analysis.

There were 1963 contigs that were classified as plasmids by geNomad. Of these, 369 contigs (18.8%) had an ambiguous classification. We carry these forward as ambiguous prokaryotic contigs. Three of the contigs were classified at the archaeal species level and 1591 were classified as bacteria at the phylum level or below.

- 2084 Bacteria
- 4 Archaea
- 151 Eukaryota
- 369 Unassigned prokaryotic sequences
- 46475 Viruses (or 27103, conservatively)
- 58894 Unclassified

Next, we added in the MMseqs2 assignments for Bacteria and Archaea. In total, MMseqs2 assigned taxonomy to 95282 contigs, so 12695 contigs were not classifiable by MMseqs2. Removing the contigs that were already accounted for by SSU analysis and geNomad, 49990 contigs were assigned taxonomy. 10822 contigs were assigned as no rank, root, or at the superkingdom level so we considered these ambiguous. That left 38902 assigned as bacterial and 312 as archaeal taxonomies.

- 40986 Bacteria
- 316 Archaea
- 151 Eukaryota
- 369 Unassigned prokaryotic sequences
- 46475 Viruses (or 27103, conservatively)
- 19680 Unclassified

Finally, we added in the Kaiju results for eukaryotic contigs. A total of 5822 contigs were assigned eukaryotic by Kaju. Cross-referencing the specific contig assignments for eukaryotes required matching the NCBI taxonomic IDs in the raw output assigned to eukaryotic taxonomy. Removing the contigs already accounted for by SSUs, plasmids, and viruses, we are left with 5231 contigs. Notably, 4888 of the total Kaiju eukaryotic contigs were classified by MMseqs2, but 178 were at no rank, 949 at root, and 2526 at the superkingdom level. Taking into account the contigs that were already assigned, there were 3223 contigs that were assigned as eukaryotic by Kaiju and ambiguous by MMseqs2, and 886 contigs assigned as eukaryotic by Kaiju and no assignment by MMseqs2 (subtracting those classified by geNomad as virus or plasmid), for a total of 4109 eukaryotic contigs.

- 40986 Bacteria
- 316 Archaea
- 4260 Eukaryota
- 369 Unassigned prokaryotic sequences
- 46475 Viruses (or 27103, conservatively)

- 15571 Unclassified

The Kaiju results add some ambiguity to the number of archeal and viral contigs. However, since the Kaiju assignments for archaea were within the same order of magnitude as the MMseqs2 results, we did not examine them closely. Kaiju assigns 23499 contigs as viruses and 25407 as unclassified, so it is tempting to say that many of these unclassified contigs are probably viral. We found that of the unclassified Kaiju contigs, 9915 were assigned as viral by geNomad, and 8458 overlapped with contigs that could not be classified by the other methods we used. Based on this information, we estimate that 33414 contigs are viral, for something more conservative than all of the geNomad results. This is closer to the other conservative estimate of 27103 viral contigs.

Combining everything, our final estimates for the taxonomy of the contigs are

- 38% Bacteria
- 0.3-0.7% Archaea
- 4% Eukaryota
- 25-43% Virus
- 14% Unclassified or ambiguous
